# “Destructive fishing” – a ubiquitously used but vague term? Usage and impacts across academic research, media, and policy

**DOI:** 10.1101/2021.05.07.443117

**Authors:** David F. Willer, Joshua I. Brian, Christina J. Derrick, Marcus Hicks, Alerick Pacay, Arlie H. McCarthy, Sophie Benbow, Holly Brooks, Carolina Hazin, Nibedita Mukherjee, Chris J McOwen, Jessica Walker, Daniel Steadman

## Abstract

The term “destructive fishing” appears in multiple international policy instruments intended to improve outcomes for marine biodiversity, coastal communities and sustainable fisheries. However, the meaning of “destructive fishing” is often vague, limiting effectiveness in policy. Therefore, in this study we systematically reviewed the use of “destructive fishing” in three record types: academic literature, media articles, and policy documents between 1976-2020. A more detailed analysis was performed on sub-sets of these records, considering the extent to which the term is characterised, geographic distribution of use, and specific impacts and practices associated with the term. We found that use of “destructive fishing” relative to the generic term ‘fisheries’ has increased since the 1990s. Records focused predominantly on fishing practices in South-eastern Asia, followed by Southern Asia and Europe. The term was characterised in detail in only 15% of records. Habitat damage and blast/poison fishing were the most associated ecological impacts and gear/practices, respectively. Bottom trawling and unspecified net fishing were regularly linked to destructive fishing. Importantly, the three record types use the term differently. Academic literature tends to specifically articulate the negative impacts, while media articles focus generally on associated gears/practices. Significant regional variation also exists in how the term is used and what phenomena it is applied to. This study provides evidence and recommendations to inform stakeholders in any future pursuit of a unified definition of “destructive fishing” to support more meaningful implementation of global sustainability goals.

## Introduction

Wild capture fisheries are a cornerstone of the global food system, drawing from enormously productive and diverse ocean ecosystems to feed billions of people. Food from the ocean currently accounts for 17% of global edible meat production (Costello et al., 2020), and marine capture and mariculture production stood at 115.2 Mt in 2020 (FAO, 2020). Sustainably managed marine fisheries have the potential to contribute to several societal needs, including ending poverty, ending hunger, decent work, reducing inequality, climate action and restoring marine biodiversity (Singh et al., 2018). Consequently, the need to tackle negative aspects of fisheries is embedded in international, regional and national policy frameworks and action plans (Singh et al., 2018). As with any policies arising from consensus-driven processes, moving from political ambitions to implementation is an enduring challenge (Armitage et al., 2020; Liuzza, 2021; Sorkar, 2020). A well-established component of this challenge is interpreting the language of global goals and their associated targets, particularly where texts of agreements, or related resolutions or measures, are legally binding for terms that are vague (King, 2017; UNESCO, 2020).

Where goals, targets and indicators have been established and gained global traction, efforts have been made to develop a more coherent, shared understanding of key words, phrases or concepts within relevant frameworks and amongst relevant stakeholder groups. In the fields of marine conservation and fisheries management, several recent examples exist of definition-setting and indicator-setting processes to aid the interpretation of terminology. These include definition-setting processes for the terms “other effective conservation measures” (a term in the UN Convention on Biological Diversity) (Alves-Pinto et al., 2021), “industrial fishing” and “levels/scales” of Marine Protected Areas (Grorud-Colvert et al., 2021), and “illegal, unregulated and unreported fishing” (Macfadyen et al., 2019). It is vital to note that these are explicitly political processes. This often means the need to satisfactorily resolve negotiations or political disputes, and reconcile collisions of divergent worldviews, interests and value systems, drives outcomes as much as – if not more so – than scientific and linguistic definitions surrounding the focal terms (J. C. Rice, 2011).

One of the most relevant global policy ambitions to fisheries is Sustainable Development Goal (SDG) 14 Life Below Water, particularly Targets 14.4 (“effectively regulate harvesting”) and 14.6 (“prohibit certain forms of fisheries subsidies”). These targets collectively refer to three problematic dimensions of fisheries: “overfishing”, “illegal, unreported and unregulated fishing (IUU)” and “destructive fishing practices”. Whilst there are established indicators to monitor progress towards ending “overfishing” and “IUU”, no such indicator exists for “destructive fishing”, limiting the effectiveness of “destructive fishing” as a policy term.

**Figure 1.**
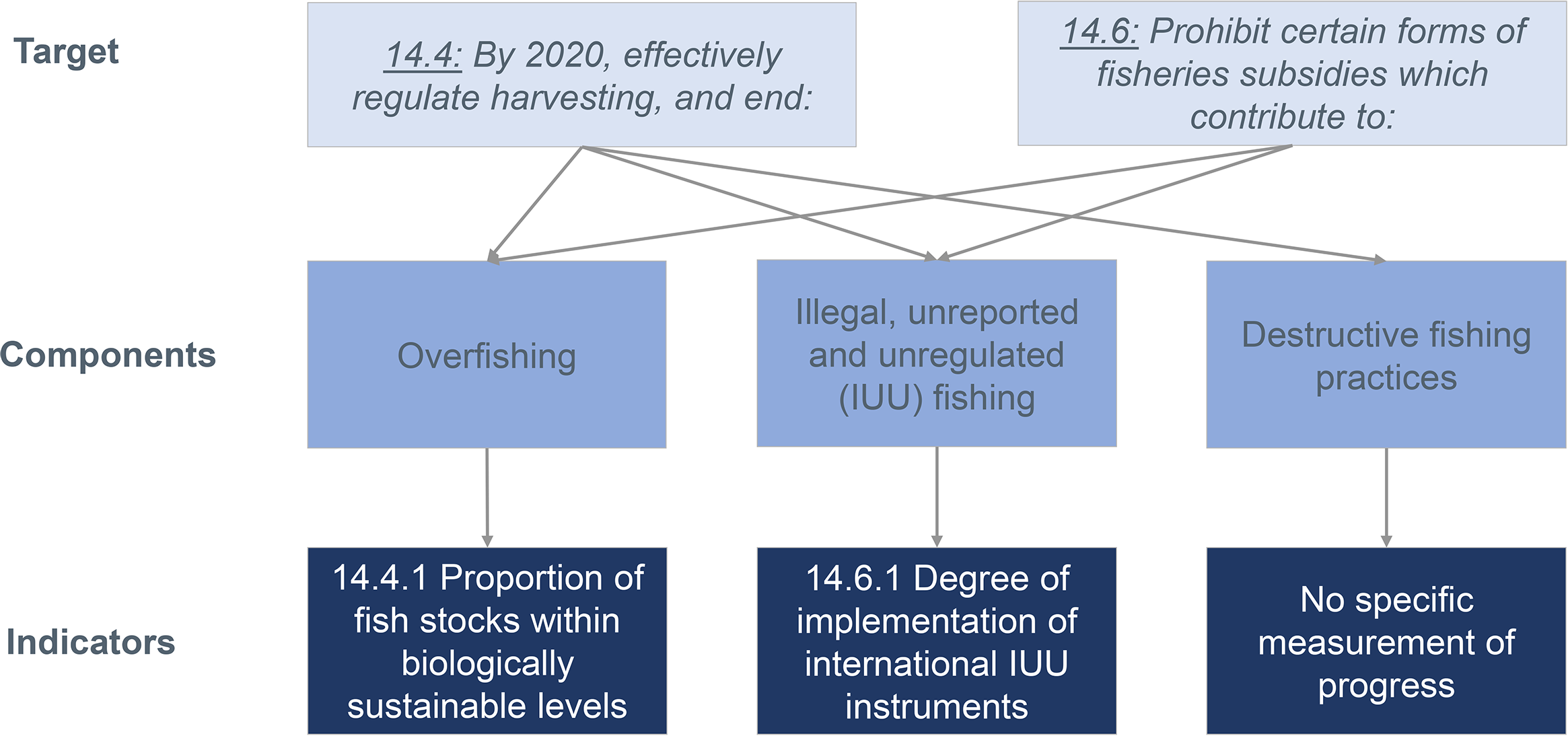
Suggested relationship between problematic dimensions of fisheries referred to in SDG Target 14.4 and 14.7 and their associated indicators.

**Table 1.** Definitions of problematic dimensions of fisheries (otherwise known as components of the overarching term “unsustainable fishing”) referred to in SDG Targets 14.4 and 14.7 (from FAO, UNEP, 2010) (FAO/UNEP, 2009)

| Component of<br>“unsustainable fishing” | Definition |
| --- | --- |
| Overfishing | “A situation in which the fishing pressure exerted on the target species is higher than the pressure theoretically required for harvesting the maximum sustainable yield (MSY), or would, if continued in the medium term, impair the population productivity” |
| IUU fishing | “Illegal, unreported and unregulated fishing”; more detailed definition in International Plan of Action to Prevent, Deter and Eliminate Illegal Unreported Unregulated Fishing |
| Destructive fishing practices | “The use of fishing gears in ways or in places such that one or more key components of an ecosystem are obliterated, devastated or ceases to be able to provide essential ecosystem functions” |

The terms “destructive fishing” and “destructive fishing practices” appear in at least five multi-lateral policy frameworks in addition to the SDGs (Figure 2, Table 2), all of which seek to “end”, or “prohibit” this problem. The intent of these suggested prohibitions encompasses supporting ecosystem recovery and sustainable resource use. The specific practices considered to be destructive also vary and include “dynamiting”, “poisoning” and “bottom trawling”, on certain habitats.

**Figure 2.**
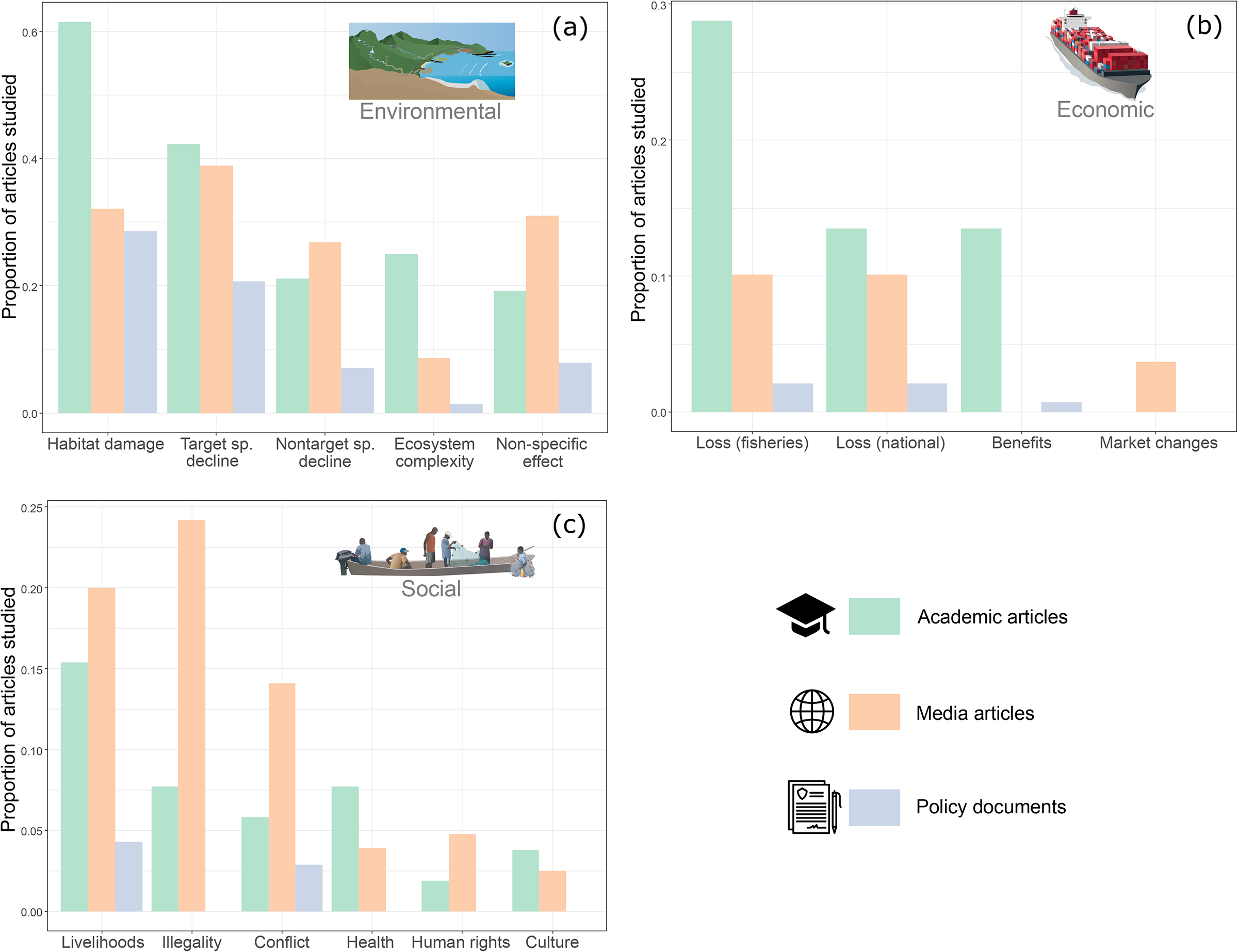
Presence of “destructive fishing” in multilateral ocean policy frameworks.

**Table 2.**
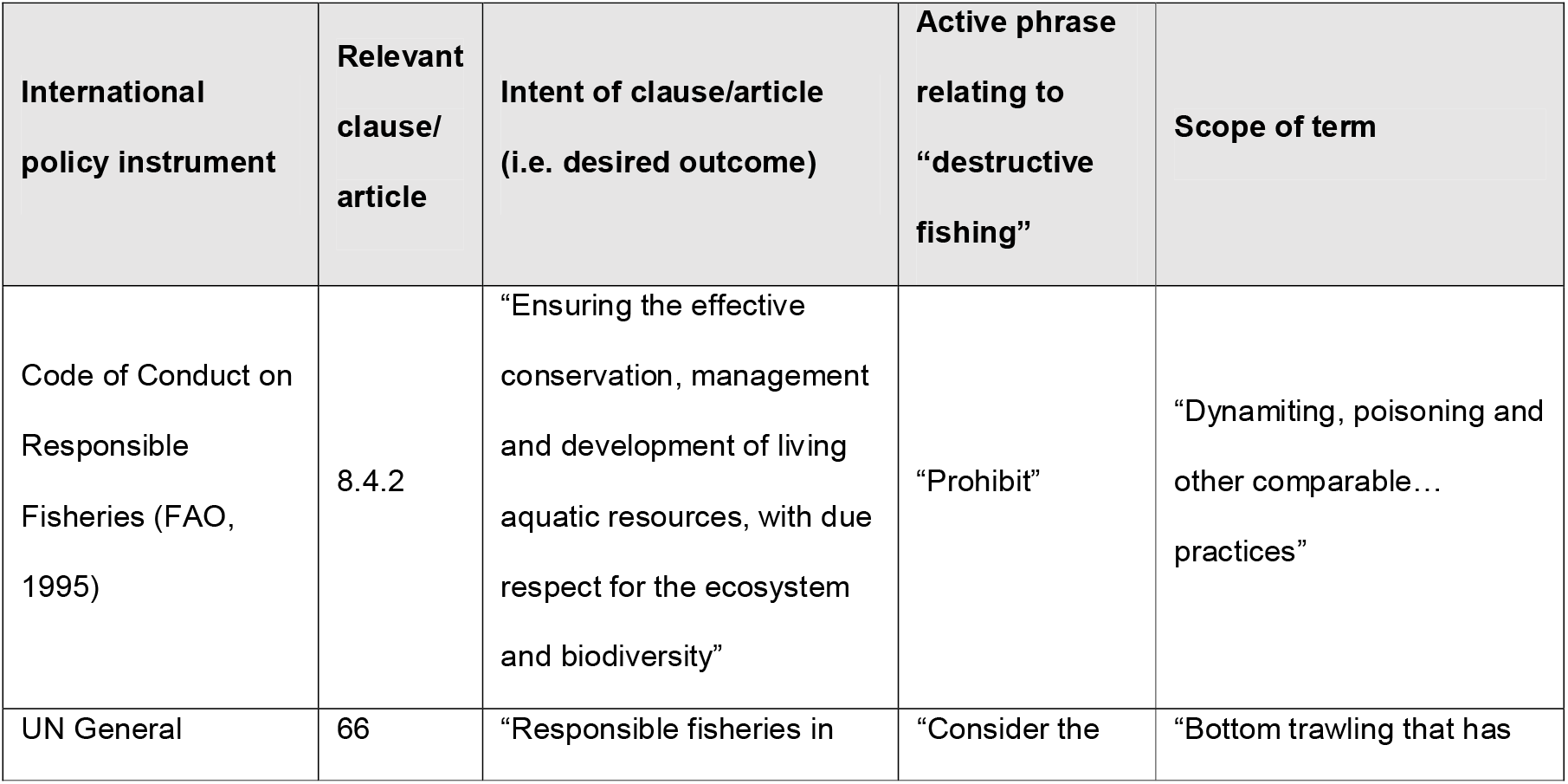
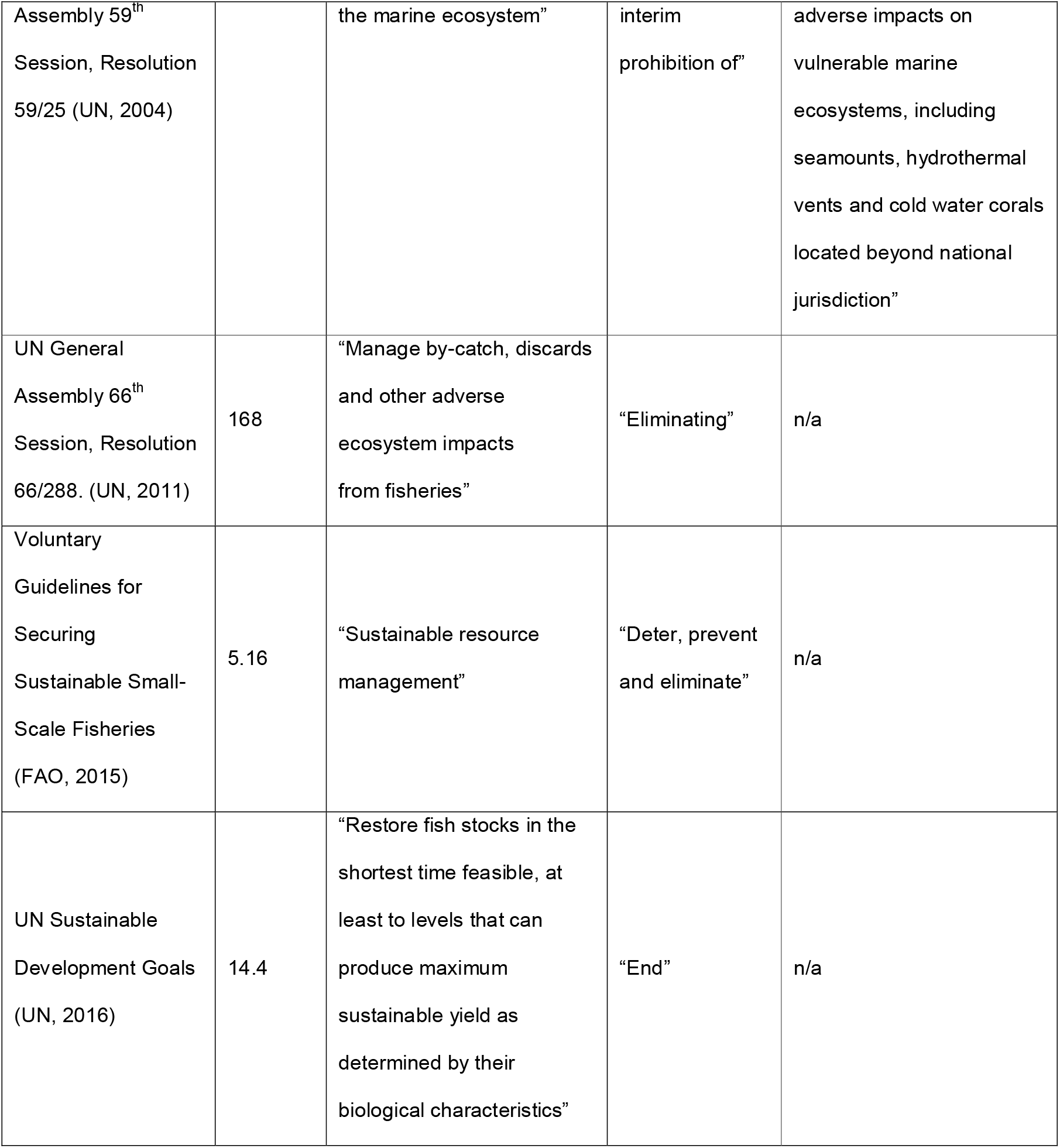
Contextual use of the term “destructive fishing” in five international policy instruments.

In a 2009 UNEP/FAO expert workshop, “destructive fishing” was described alongside “IUU” and “overfishing” as a sub-component of the term “unsustainable fishing” (FAO/UNEP, 2009). In this workshop “destructive fishing” was described as “the use of fishing gears in ways and places…[such that]…one or more ecosystem components are obliterated, devastated or ceases to be able to provide essential ecosystem functions” (Table 1). This description has not been formally ratified as an internationally agreed definition. The description also states that “only a very small number of fishing gears or fishing methods are recognized as inherently ‘destructive’ wherever and however they are used, the primary examples being explosives and synthetic toxins. In the absence of any formal agreement regarding the term, the classification of a gear or practice as destructive is a policy choice related to pre-set objectives and consistent with national and international law” (page 9 in FAO/ENEP, 2009).

This summary alludes to an unresolved tension of values/worldviews around the discussion of “destructive fishing” that (J. C. Rice, 2011) expands on, noting that FAO and UNEP experts in the cited workshop differed significantly in their approaches to synthesising the evidence they presented in support of an improved definition. Recognising historical tensions and barriers to meaningful progress is vital to inform any future attempt to improved shared understanding of “destructive fishing”. In particular, tensions have been around 1. Whether scientists and/or experts should explicitly direct policy-makers as to which practices are “categorically harmful or acceptable” and 2. Divergence in expert opinion as to the relative inherent destructiveness of specific fishing gears. Interestingly, while the workshop itself appears to have been seen as productive, the subsequent involvement of individuals beyond original participants fostered further intersectoral tensions resulting in attempts to accommodate an unwieldly number of perspectives, thereby hindering the imperative to make the term more specific (J. C. Rice, pers. comm).

In addition to its presence in a multitude of policy fora, the term “destructive fishing” frequently occurs in popular ocean discourse and is used by a range of other societal actors (e.g., academic literature, NGO campaigns, media coverage) (Ochave, 2016; Odvek, 2021). This is perhaps not surprising given that the word “destructive” is one with strongly negative associations. When such words are used in a public discourse without context or in a vague manner, they can drive intersectoral and political polarisation (Cap, 2017). We hypothesise that “destructive fishing” and “destructive fishing practices” have the potential to become and indeed are predominantly utilised as “quasi-concepts”. The terms as they stand are, in effect, “flexible enough to allow the meanderings and necessities of political action from day to day” (Bernard, 1999). Therefore, it is critical to assess the use of “destructive fishing” across the areas it is used, which includes the media and academia as well as in policy documents, to develop a definition which is accurate and useful in informing global policy.

We think that a unified definition of “destructive fishing” would produce benefits for sustainable fisheries management and marine conservation by reducing intersectoral polarisation on the definition, increasing alignment of political objectives and influencing “on the water” implementation of those objectives. Recognising the political nature of definition-setting and the challenges faced by previous attempts to gain consensus around the term, we seek to justify why a unified definition is required and provide a comprehensive overview of current usage of the term to inform progress towards an acceptable and practical definition. Through systematically reviewing the term’s usage in English-language academic articles, media articles and policy documents, we will attempt to explain the drivers of its use and consider the consequences of leaving the term undefined. Specifically, we aim to address the following questions:

1. Has the term’s relative usage over time increased?
2. How is its English-language use distributed geographically?
3. How often is the term explicitly characterised or explained?
4. What specific impacts (environmental, social and/or economic) are associated with the term’s usage?
5. What is the scope of the term’s usage in relation to specific “practices/gear types” that are referred to as “destructive”?

## Methods

### Data extraction

References to destructive fishing were extracted for academic literature, policy documents, and media articles (Table 3), in the English language only. All databases were searched using the term “destructive fishing”, and records were extracted if the term was found in the in the title, abstract/introduction and/or body text of records. Academic literature was extracted from the Scopus (Elsevier) database on the 1^st^ of March 2021; this database contains records from approximately 35,000 journals in the life, social, physical and health sciences. Policy documents were extracted from the FAOLEX (United Nations) database on the 4^th^ of October 2021; this database is administered by the Food and Agriculture Organisation, and is one of the largest online repositories of national laws, regulations and policies on food, agriculture and natural resources management. Media articles were extracted from the Factiva (Dow-Jones) database on the 5^th^ of August 2021; this database combines over 30,000 newspaper, website and online news sources. All articles up until the search date were included, so the three searches cover slightly different time periods. In addition, the date of earliest record varied between the three databases; we controlled for this discrepancy where appropriate (see below). To justify our selection of English language only content, we briefly screened all three databases for the term in Spanish (“pesca destructiva”). This screening returned 3.5% (n=5) of policy and legal documents, 5.8% (n=274) of media articles and no academic articles.

Subsets of the English language extracts were selected for more detailed analysis and characterisation (see below). We selected a subset of records from each database because the volume of total records (particularly of media articles) would mean detailed analysis would be prohibitively time-consuming. For policy and academic records, we analysed those most likely to be explicitly concerned with destructive fishing, rather than just mentioning it in passing (see “Rationale for selecting analytical subset” in Table 3). For media articles, no reasonable criteria existed and so a random sample was chosen, with the sample size determined by the number of records required to be statistically representative at the 95% confidence level. To ensure the difference in 2021 coverage did not bias results, we excluded 2021 records from all subsets.

**Table 3.**
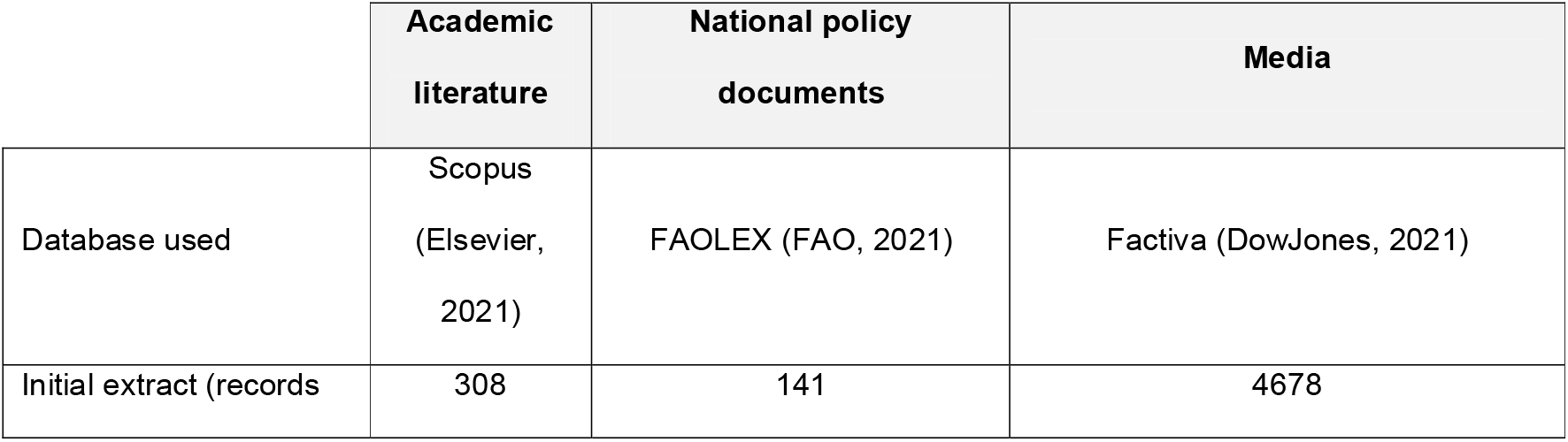
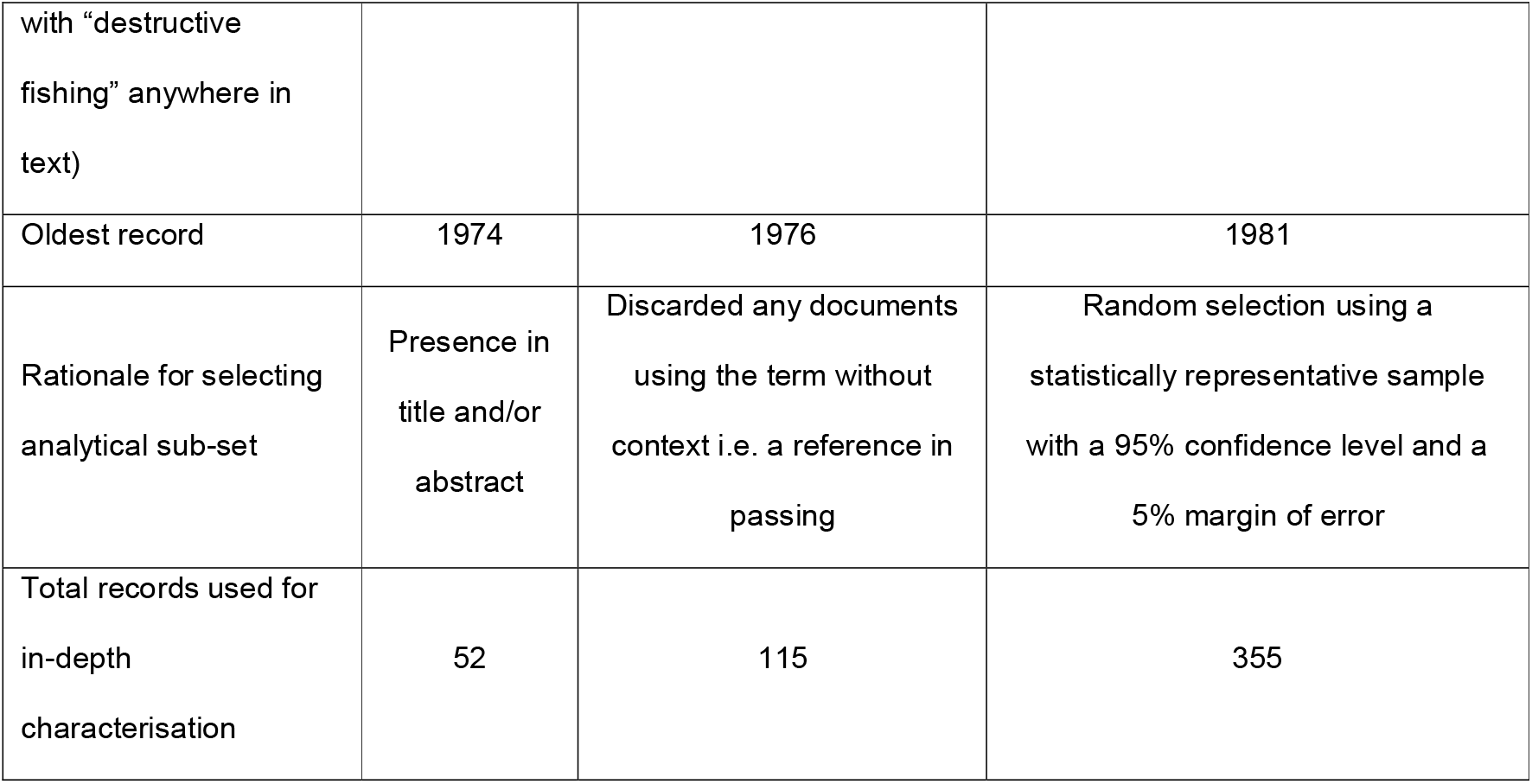
Description of databases used and sampling methods.

### Temporal analysis

To calculate the rate of publications concerning “destructive fishing” (i.e., temporal publication trends), while accounting for the increased background rate of publications generally, we first took the total number of relevant records extracted from each database (“Initial extract” in Table 3), and calculated the number of records per year. Then, we searched each database again using the term “fishery OR fisheries”, and extracted the total number of records per year. For each database, we then divided the total number of “destructive fishing” records per year from the total number of “fishery OR fisheries” records per year, to generate a “relative publication rate” metric that tracked terminological use relative to all content in this topic domain. In other words, this metric gives the proportion per year of all records to do with fish and fisheries that mention destructive fishing, and thus provides a fair estimate of the rate of increase or decrease in interest in destructive fishing while accounting for background publication trends. To allow comparison among record types on the same axis scale, we limited the years to be between 1980 and 2020 for this analysis; <5% of records were before 1980 in all of the databases.

### Geographical distribution

For the subset of records chosen for in-depth analysis, we determined the geographical distribution of records by recording the focal country/region in the academic literature, media articles and policy documents, which we used to plot the overall distributions of regions of interest for each record type. In addition, if the information was available, we also recorded its geographical origin (i.e. the home country of its host publication; see Dataset S1). Countries were subsequently assigned to a region based on either its “sub-region name” or “intermediate region name” according to (UNSD, 2021). Records that were global in scope or did not specify a focal country were discarded for this analysis only. In total, the focus region could be determined for 471 out of 522 records (36/52 academic, 113/115 policy, 322/355 media).

### Characterisations and associated impacts and practices

For the subset of records that were chosen for in-depth analysis (focal articles), information relating to the use of the term “destructive fishing” was characterised in three ways. First, we recorded whether the record provided a characterisation or explanation specifically of the term destructive fishing; if so, we recorded the characterisation and noted its key properties. We then undertook additional quantitative analysis to note where the term was associated with specifically named negative impacts and practices/gears (i.e. that were inferred as being destructive through contextual use), described in the next two paragraphs. We have provided information for each record in the subset, in addition to their geographical distribution and how we coded their listed impacts and gear types, in Dataset S1.

Second, we carried out a form of iterative analysis, following (Srivastava & Hopwood, 2009), to document the impacts associated with the use of the term, as described by our records. We began by reading a small selection of the focal articles while asking the question ‘what specific impacts do these authors link with destructive fishing” and writing individual entries for the specific impacts listed in each. During this initial process it became clear that each entry belonged in one of three thematically grouped categories: environmental, social or economic changes. We therefore established these categories as overarching themes, and then arranged individual entries into subcategories we developed that nested within each theme (e.g. “destroying coral reefs” and “ripping up seagrass” would both be classed in the ‘Habitat destruction’ subcategory in the environmental theme). These subcategories were then further developed in iterative fashion, by reading the rest of the focal articles and placing described impacts into the different subcategories, adjusting or adding subcategories where necessary. This ensured that the nuance in the impacts listed had been captured by our themes and subcategories, thus reconciling the relationship between what the data said and what we wanted to know (Srivastava & Hopwood, 2009).This process led to six subcategories in the social theme, five in the environmental theme and four in the economic theme. To ensure both consistency and reliability in categorisation, the iterative analysis was carried out by two authors (A.P. and J.W.) who developed the subcategories, checked each other’s categorisations and agreed upon the final classifications.

Third, we explored the specific practices/gear types that were associated with the term “destructive fishing.” Based on the expert knowledge of the author group (i.e. in contrast to the content analysis above) we *a priori* identified three categories that practices could fall into: 1. The use of a specific fishing gear (e.g., beach seines), 2. The use of an auxiliary device/gear component (e.g. lights on catching devices) or 3. Other practices/fishing-associated behaviours (e.g. “trash fishing”). Where references to practices were sufficiently detailed in the reference to specific gears and/or auxiliary devices, we used the classification system in (He et al., 2021), which is an objective, multi-lingual lexicon of fishing technology developed by FAO gear technologists. From our records, we identified 40 separate fishing practices associated with the term “destructive fishing” across the three content types; 24 identifiable fishing gears, five auxiliary fishing devices and 11 other fishing practices/fishing associated behaviours. We calculated the proportion of references to each fishing practice and derived a mean proportion across all content types; practices with a mean proportion of less than 2% were discarded from further analysis.

Finally, we carried out an additional cross-table analysis to break the impacts and gears down by record type (academic, media or policy) and geographic distribution (at a continental scale, with Central and South America being combined), to determine if there were different trends between continents for listed gear types and impacts. To account for the different numbers of total records between continents, we calculated the percentage of records within each document type and continent that mention a specific impact or gear type, excluding those groups where there were <3 total records (Table S1 Gears, Table S2 Impacts).

## Results

The study reveals a large increase in the use of the term “destructive fishing” in academic literature, media articles, and policy documents over the past four decades, even while accounting for increases in the background publication rate (Figure 3). It is notable that the highest publication rate values for academic literature and policy documents occur after the 2015 Agenda for Sustainable Development, the process that created the SDGs. Similar sized spikes occur in academic records in the mid 1990s after the development of the Code of Conduct for Responsible Fisheries in 1995; the fact that these spikes are not reflected in national policy records logically suggests a time lag between the ratification of global instruments and their inclusion in national decision-making frameworks, although it is notable that policy documents have seen the sharpest overall increase in usage rate.

Media usage of the term has risen the least sharply of the three content types and its only notable spike comes after UN Resolution 59/25 (on deep-sea fisheries) in 2004, which explicitly refers to case-specific instances of high seas bottom trawling as “destructive fishing” (Table 2). This initial adoption by the UN General Assembly to introduce more precaution into how high seas fisheries are managed was followed by an intense period of political campaigning (between 2005-2008) for a moratorium on high seas bottom trawling (Carmine et al., 2020).

**Figure 3.**
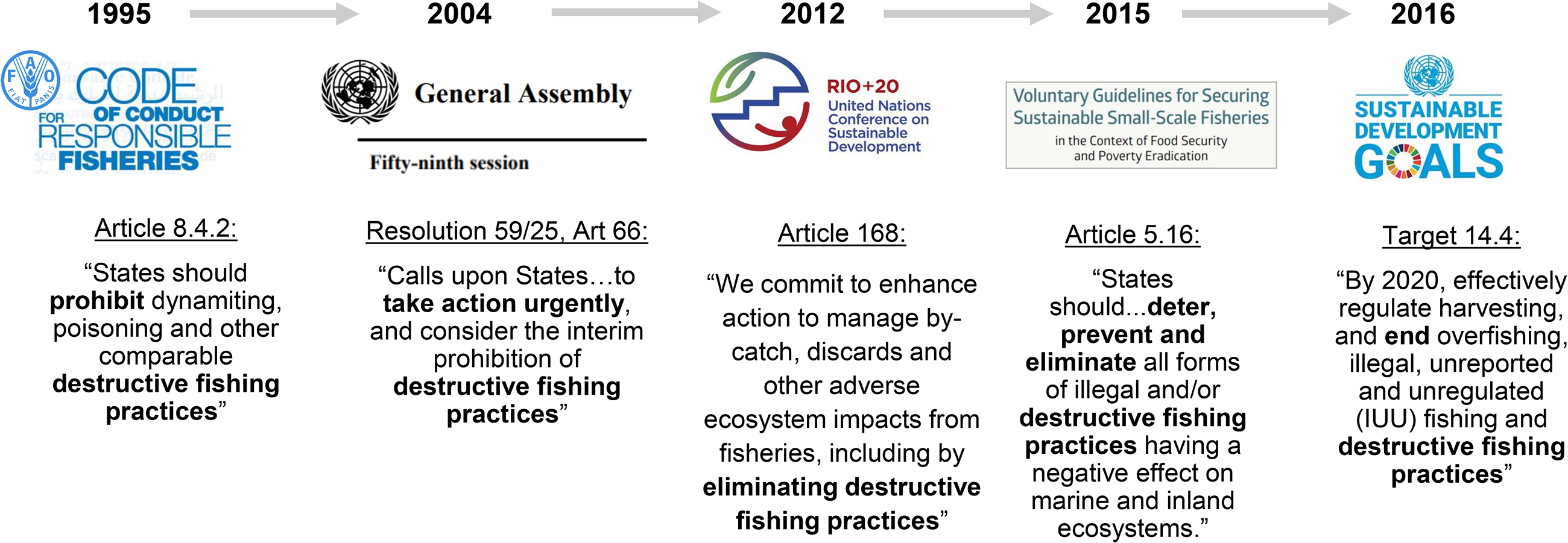
Change in frequency articles focusing on destructive fishing over time. Articles are in academic literature (green line); media articles (orange line); and policy (blue line). The article frequency rate is adjusted to account for the background rate of publications on fisheries; see Methods. Vertical lines in (labelled a - i) indicate significant global policy mechanisms that impact fisheries management and conservation

Most records across academia, media, and policy relating to “destructive fishing” focus on practices in South-eastern Asia (61%, 38% and 23% of academic, media, and policy respectively) (Figure 4). This is followed by Southern/Western Asia, Oceania, and East/South Africa which each represented at least 5% of the academic, media, and policy records. Some of these differences are striking: for example, there were no academic articles focusing on destructive fishing in the Americas, and very few in Europe, in contrast to low- and middle-income regions in the tropics which were disproportionately represented in the academic literature.

There were also clear discrepancies between the distribution of record types between geographic regions. For example, North America and Europe had the greatest proportion of their records coming from the media, while Southeast Asia and Southeast Africa had the greatest proportion of their records coming from academic articles. Oceania, Central America and West/North Africa had the greatest proportion of their records coming from policy documents (Figure 4).

**Figure 4.**
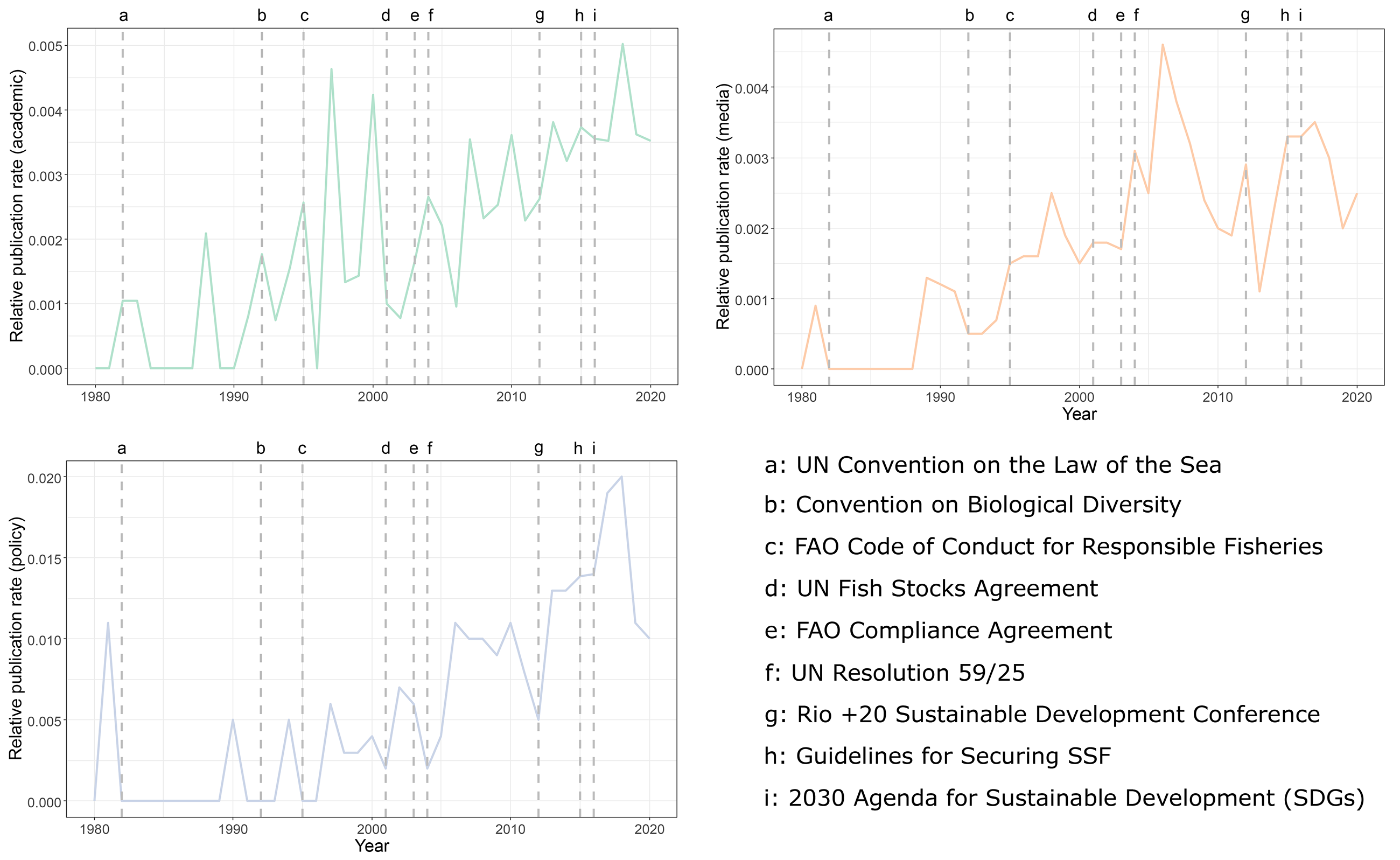
Map indicating the percentage of academic, media, and policy focal articles that focus on each global region. Alternating shades of grey are used to represent each global region. Percentages were rounded up to the nearest whole percent, to enable records at <0.5% to be visualised, therefore some totals slightly exceed 100%.

The term “destructive fishing” was only defined and/or characterised in 13% of the academic literature (n=7, out of 52), 19% of policy documents (n=22, out of 115) and 14% of media articles (n=50, out of 355). A sub-set of those characterisations are presented in Table 4.

**Table 4.**
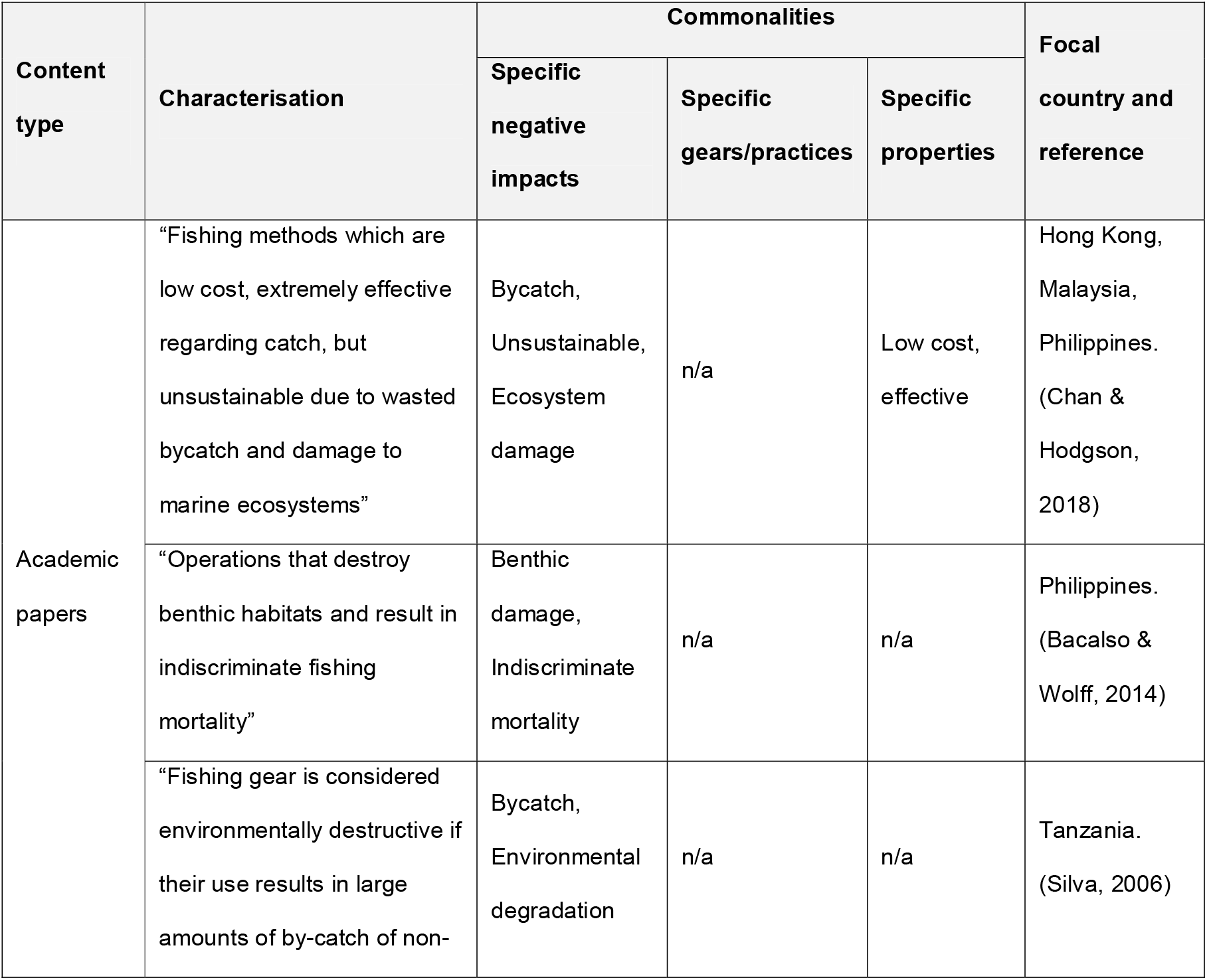
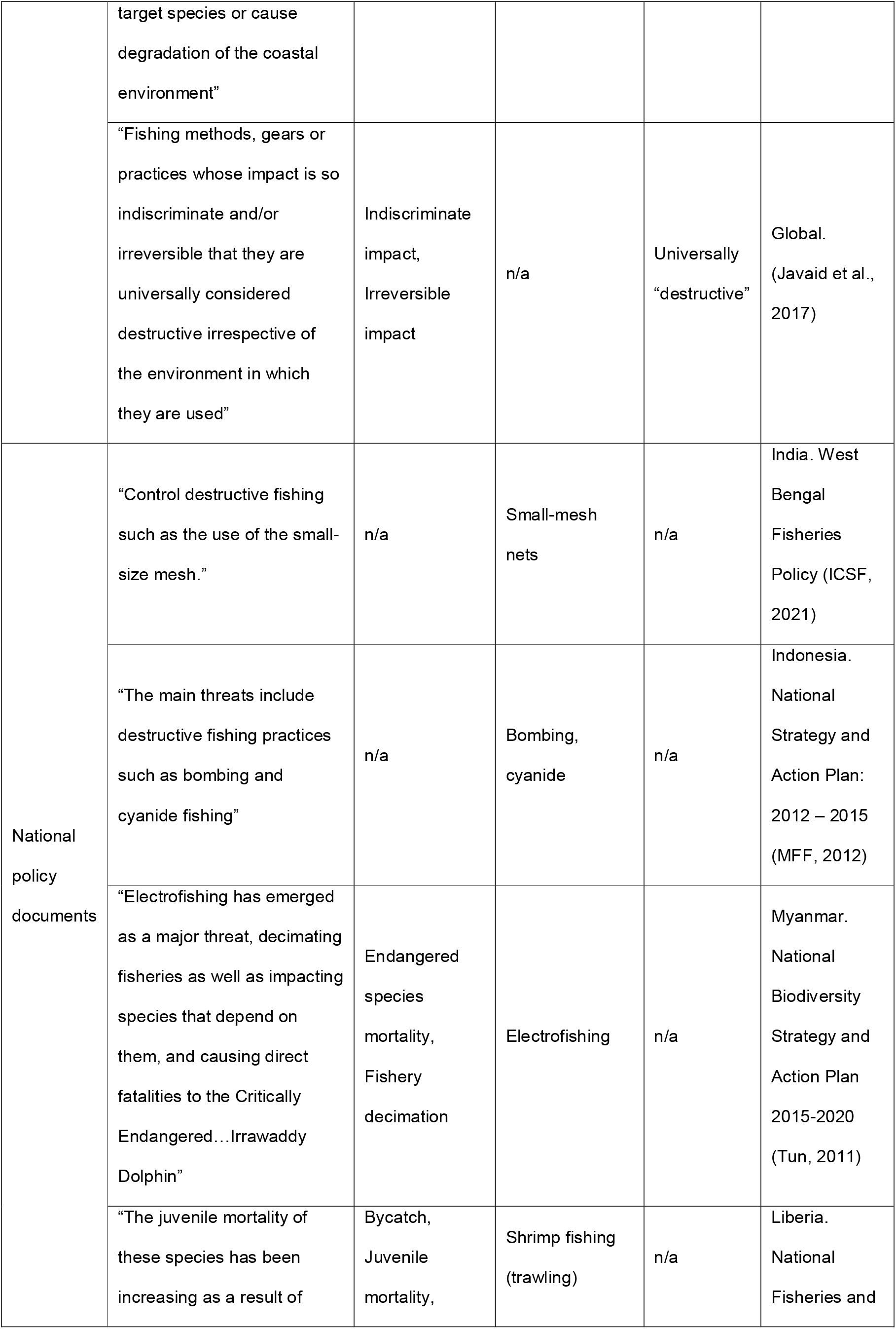
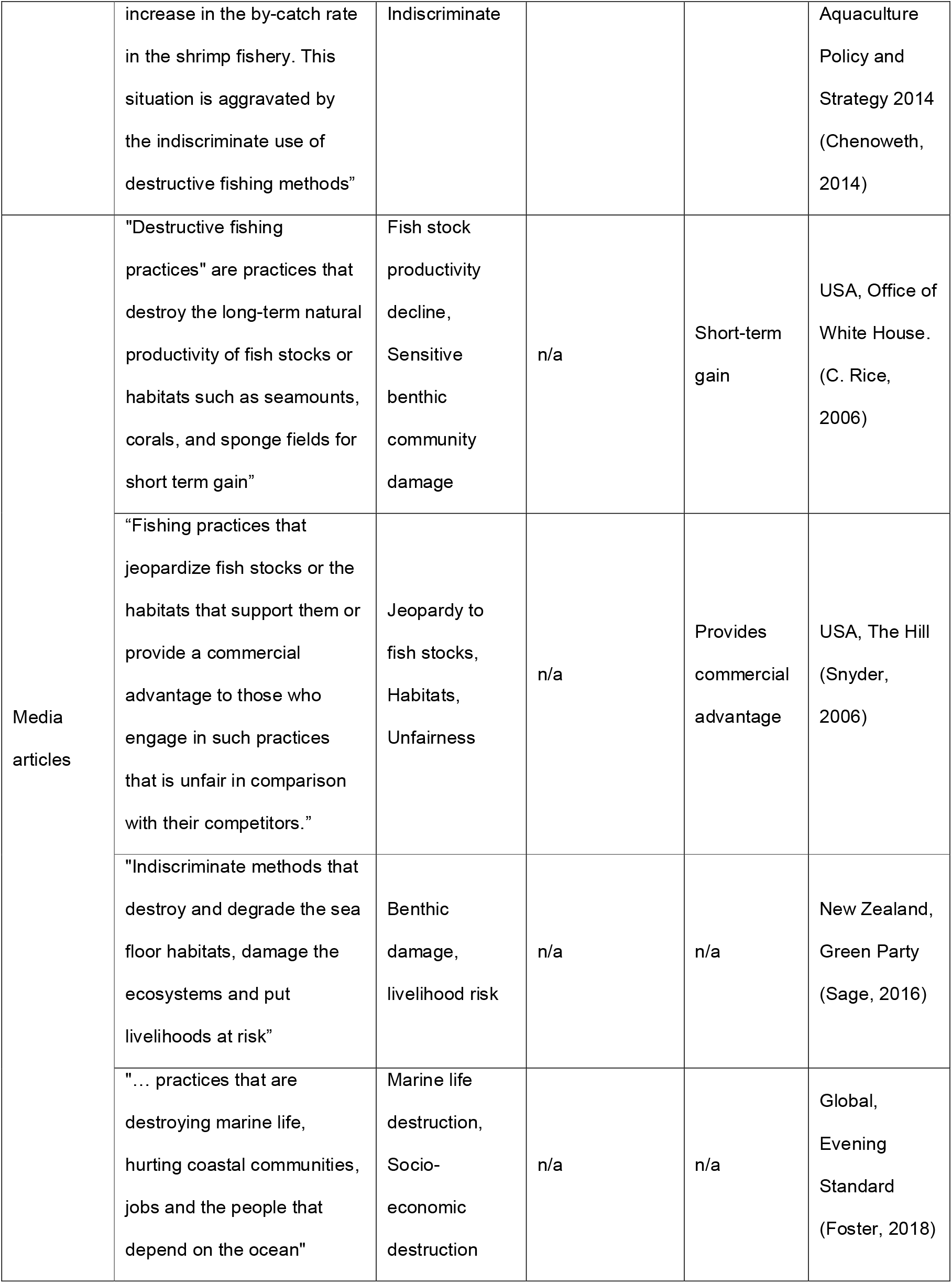
Examples of the term “destructive fishing” and commonalities between examples from across record types. See Dataset S1 for full details.

Across the records, environmental, economic, and social impacts associated with “destructive fishing” were identified (Figure 5). The proportion of records considering all of the impacts was highest in academia, with almost all (94%) of the literature focusing on the impacts, compared to 85% of media articles, and 61% of policy documents. Environmental impacts were the most reported type of impacts across all three record types (Figure 5a); predominantly habitat damage, closely followed by target-species population decline, with the exception being a greater prevalence of target-species decline in media articles.

The media had the greatest focus on social impacts (30% of all reported impacts, compared to 16% in academic literature and 9% in policy), with illegality and damage to livelihoods dominating (Figure 5b). The academic literature reported economic harm the most frequently (21% of all impacts, compared to 10% in the media, and 6% of policy), with loss to local fisheries and fishers’ livelihoods being the most prevalent concern. The short-term economic benefits of destructive fishing to the individual fisher were raised in the academic literature and infrequently in the policy documents, but were not reported in the media articles. In contrast to the academic literature and media articles, policy documents focused more on environmental impacts than broader social and economic harm.

There were also some clear differences in the distribution of impacts listed between geographic regions. For example, within media articles, the economic losses to fisheries in the private sector caused by destructive fishing were heavily emphasised in articles from Africa and Asia, with little mention in media articles from other continents (Table S2). In contrast, the environmental impacts of habitat damage and non-target species decline were disproportionately mentioned in media articles from Europe and North America, with a much lower rate of mention in Africa and Asia; whilst the decline of target species was mentioned fairly evenly across media articles from all continents (Table S2). This was in striking contrast to the policy documents, where the impact of the decline of target species was not mentioned in any policy document from Europe or North America.

**Figure 5.**
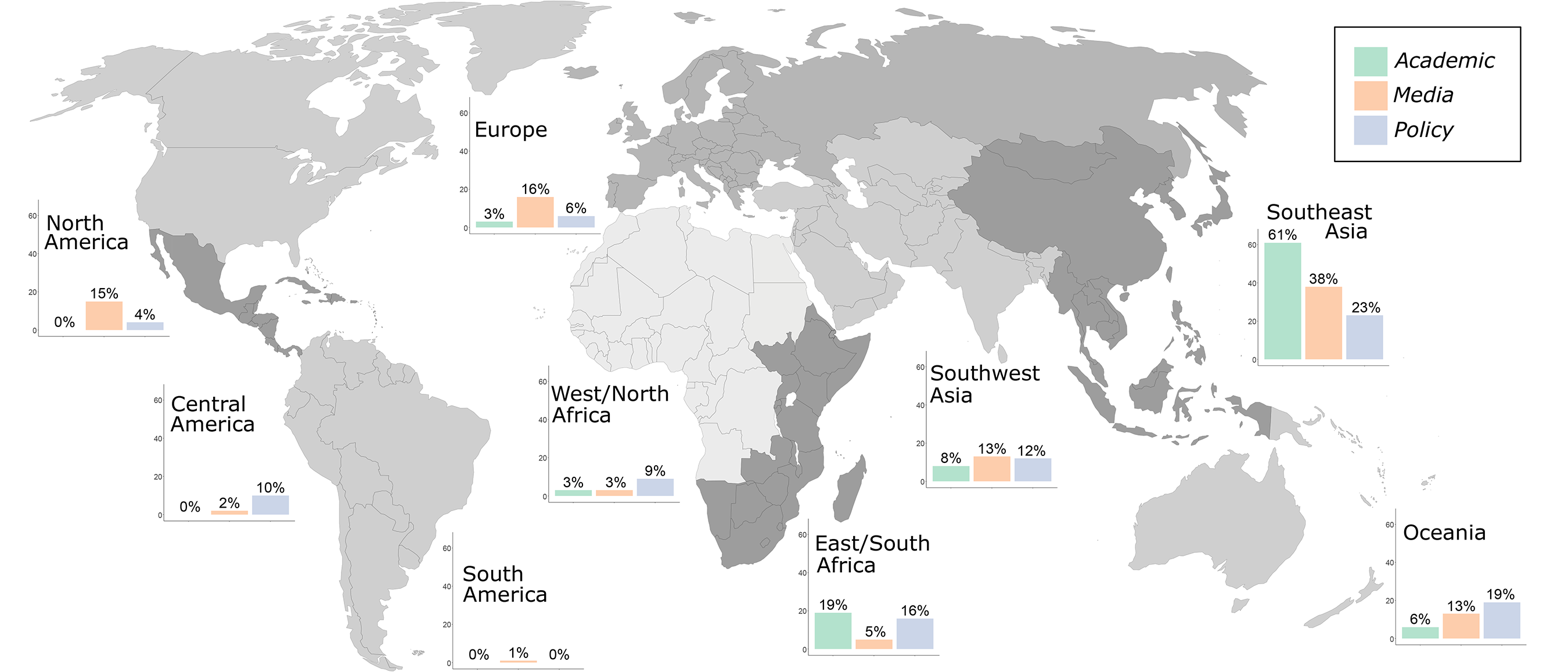
Bar charts showing the ecological, economic and social impacts of destructive fishing for each content type. Ecological, economic, and social themes are grouped into key subcategories; see Methods for the process used to define these groups. Note that there was uneven emphasis on the three themes, with environmental impacts being more widely discussed than economic or social impacts, and so each y-axis is on a different scale. Main coloured images: Tracey Saxby, Integration and Application Network (ian.umces.edu/media-library). Small icons from the Noun Project (website icon, Syawaluddin; policy document icon, iconixar; academic icon, general Noun project).

94% of the academic literature (n=49), 49% (n=56) of policy documents and 56% of media articles (n=198) mentioned at least one fishing practice (Table 5). Of the 23 practices that occurred at >2% frequency, only four had an overall proportion of references above 10%; “Blast fishing” (51%) and “Poison fishing” (43%), then “Bottom trawls” (27%) and “Nets, unspecified” (15%). There was more emphasis on the first two practices in academic literature and more emphasis on the third in media articles with nets showing a more even spread.

The distribution of gear mentions was unevenly distributed among continents within each record type. The distribution was not consistent between record types: for example, 22% of media articles from Oceania mention purse seines (no other continent had more than 4% of media articles mentioning this) (Table S1); but then Oceania had no policy documents at all mentioning purse seine, suggesting a disjunction between the emphasis of destructive fishing placed by media and policy (Table S1). Other clear areas of potential concern for management for specific geographic regions could be identified: for example, 38% of North American policy documents mentioned set gillnets, while no other region had more than 4% of policy documents mentioning this gear type; in contrast, many media articles from Africa and Asia mentioned nets while few media articles from other continents did (Table S1). We also detected the potential importance of specificity in defining gears: for example, ‘trawls’ generally were most mentioned in media and policy from Africa and Asia, while ‘bottom trawls’ were mentioned most in media and policy from North America and Europe.

**Table 5.**
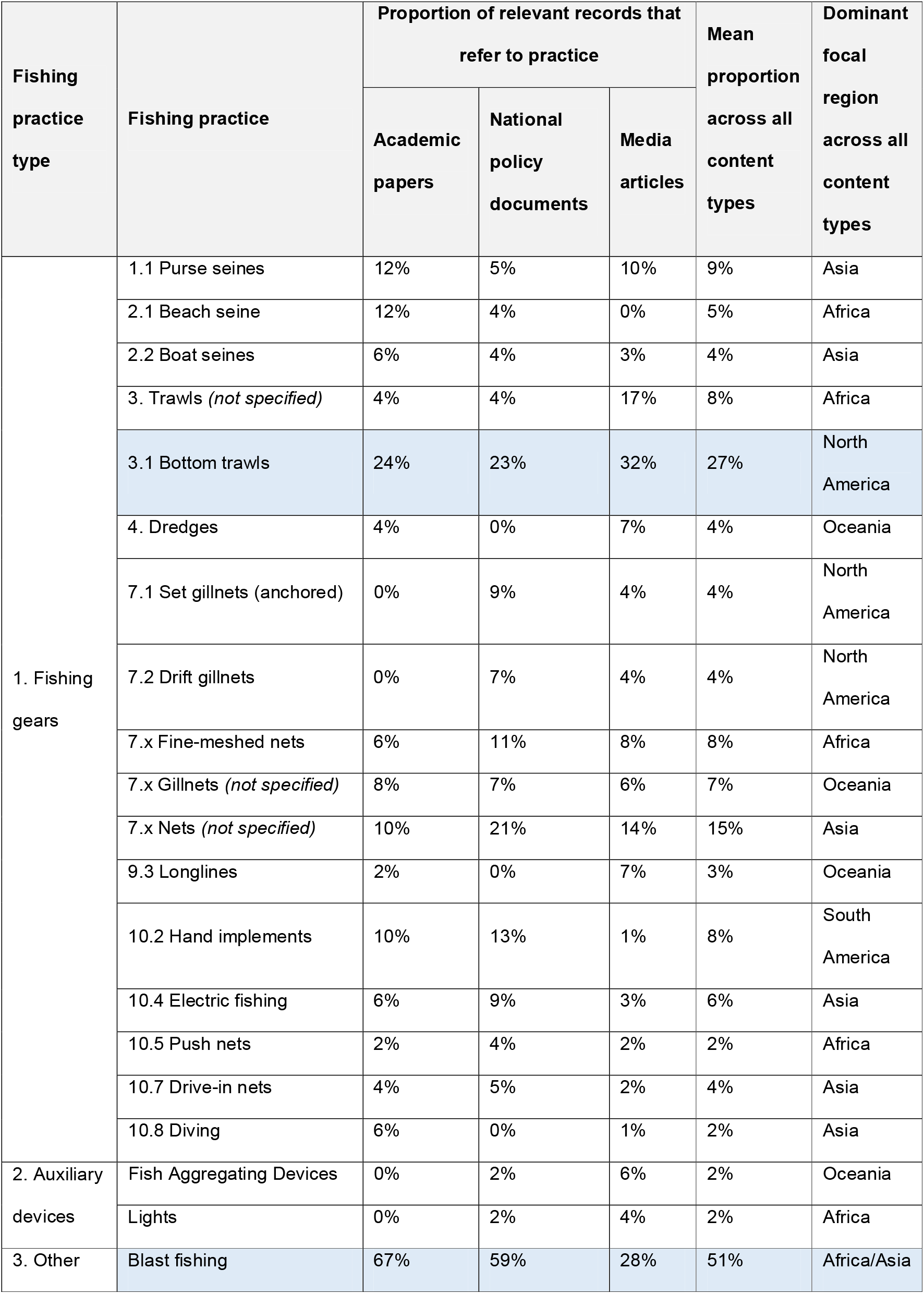
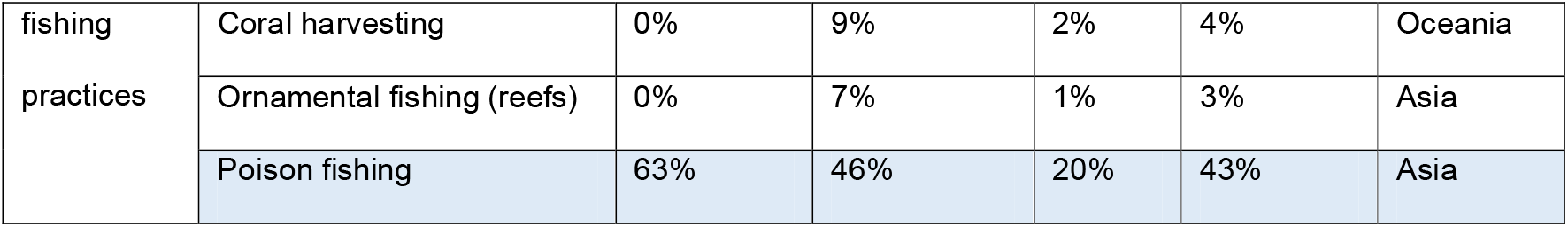
Fishing practices associated with the term “destructive fishing” and proportions of reference to each practice across record types. “Fishing gears” ordered by classification in (He et al., 2021). “Auxiliary devices” and “Other fishing practices” ordered alphabetically. Practices with mean proportion of references above 25% highlighted in blue.

## Discussion

Our results illustrate that “destructive fishing” means different things to academics, media producers and policy-makers in different parts of the world, and that moving towards a shared understanding of “destructive fishing” will require reconciling a set of contrasting yet potentially equally valid approaches to the term.

Our study shows that in the three record types of academic literature, media articles and policy documents, the relative usage of the term “destructive fishing” has increased over time. We found that its English-language use is geographically biased towards South-eastern Asia. We found only a minority of specific characterisations in each of the three record types; at 19%, policy documents had the highest proportion. We much more frequently identified negative impacts and gear/practices associated with the term’s usage (i.e. that were inferred as being destructive through contextual use). Environmental impacts – particularly habitat damage – were the term’s most consistently associated impacts and the use of explosives and poisons to fish were the most commonly associated gears/practices, with the very separate gear/practice of bottom trawling also central to the term’s usage.

Building off these findings, we now acknowledge limitations of our study, summarise why we believe this term has been (and continues to be) subject to vague usage, consider the consequences of leaving the term undefined and offer recommendations for the future pursuit of a unified definition of the term.

### Research limitations

The authors acknowledge that our analysis was limited by several factors. First, while we did screen for the term in one additional language (Spanish – see Methods) and found minimal additional references to the term, there would be value in additional consideration of even those limited references. In particular, this may explain the unusual trend of no academic articles concerning destructive fishing in Central and South America (Figure 4): they may have been absent in our English-language search because they were written in Spanish, and absent from our Spanish-language search because (to our knowledge) Scopus does not specifically index Spanish sources, in contrast to the FAOLEX and Factiva databases. While beyond the scope of the current manuscript, we highlight the need to consider the possible evidence base available in other languages when moving forward with a wider destructive fishing discourse (Amano et al., 2016). Second, we acknowledge that trend formation (i.e. the underlying drivers of why a concept emerges and becomes significant) is more complex than the basic terminological history we have been able to present. It is likely that the political, scientific and popular discourses around this term are confounding variables that influence one another in explaining the term’s usage patterns, rather than discrete factors. Finally, while we attempted to consider the differing motivations and mandates of the three record types we drew our data from, we acknowledge contrasts in how language is used in these distinct realms of discourse. In particular, we recognise the need for more analysis of ideological bias, sentiment and positionality in further explaining why and how “destructive fishing” is prioritised in these records. Nonetheless, our results offer valuable insights from which we can consider the consequences of using the term vaguely and form recommendations for future work.

### Why is the term “destructive fishing” used vaguely?

Through considering its usage in multi-lateral policy instruments (Table 2) from the FAO Code of Conduct for Responsible Fisheries in 1995 (FAO, 1995) onwards, we see that there is a consistent call for states to “end” or “prohibit” destructive fishing. However, the intent of these measures or the negative impacts they are trying to prevent is often vague, therefore the scope of the measures or specific practices they direct states to end is often absent. By confirming that usage across our three content types is also vague, we demonstrate the need for a revised definition-setting process for the term “destructive fishing”, building on past efforts to derive a unified definition (FAO/UNEP, 2009; J. C. Rice, 2011).

When considering examples of the term’s associated negative impacts (Figure 5), we found that specific negative impacts were articulated most commonly by academic literature and least commonly by policy documents. This suggests that scientific research is the most likely record type to try to identify a specific effect around the term “destructive”. This finding is complicated by the abundance and diversity of associated impacts across all three record types. While environmental harms such as “habitat damage” and social harms such as “damage to livelihoods” were relatively consistent, other identified impacts pointed towards “destructive fishing” overlapping with other, more-defined, problematic dimensions of fisheries (e.g. “target species decline” and “overfishing”, “illegality” and “IUU”, “unsustainable fishing”). Separating what is “destructive” from these more established concepts is vital in ensuring clarity in the term’s future usage.

Regarding the term’s associated gears/practices (Table 5), we found that specific gears/practices were identified most commonly by academic literature and least commonly by media articles. This finding broadly supports the notion that scientific research is more likely to attempt to identify a specific action as “destructive” than a media article is. There is also different emphasis placed on both gears and impacts between policy and media *within* each continent (Tables S1, S2). It suggests the importance of different concepts, as deemed by policy-makers, are poorly reflected in the media, who may be more driven to generate more general interest in destructive fishing. More generally, the very different emphasis on impacts among record types (Figure 5) indicates that different stakeholders have very different interpretations on how the term should be used, providing a key reason why the term, as it stands, is so nebulous.

The focus on negative impacts and gear/practices associated with “destructive fishing” are perhaps also an explanation of why the term is used vaguely. These impacts and gears/practices are the most common markers associated with the term’s written use and it is discrepancy around these markers that inhibited past attempts to define the term (J. C. Rice, 2011). Selecting simple impacts and gears/practices enormously simplifies the complexities outlined in (FAO/UNEP, 2009) and the range of spatial, temporal and regional dimensions of what may constitute “destructive fishing” as well as what constitutes a “practice”. Our findings also emphasise that discourse around this term partly driven by political and value-oriented discussions of “which fishing gears cause which environmental harms”. This is instructive in explaining why the term is vague given that the discourse generally remains unresolved and polarised.

We emphasise that “destructive fishing” means different things to academics, media producers and policy-makers in different parts of the world, and that a shared understanding of “destructive fishing” requires reconciling a set of contrasting yet potentially equally valid approaches to the term. We see this trend emerge in three specific ways in our results. First, different gears are emphasised by the records of different continents across all of academia, media and policy (Table 5, Table S1), suggesting that different parts of the world may be subject to different destructive practices (or may differentially ascribe destructive properties to a practice). Therefore, a ubiquitous approach to “destructive fishing” may benefit more from identifying *outcomes* as destructive, rather than specific gears, which vary in usage and impact throughout the world. Second, we also see a different focus in impacts between continents (Table S2), which may reflect the differing importance in fishing more broadly. In particular, media records from Africa and Asia were particularly concerned about “destructive fishing” causing a loss in fisheries income, and policy records were concerned with the decline of target species. In contrast, media from Europe and North America were more concerned with habitat damage and non-target species decline, and policy documents were not concerned at all with target species decline (Table S2). We suggest that this may reflect the increased importance of small-scale and subsistence fisheries in low/middle income tropical regions relative to high income temperate regions. The recognition that fisheries (on the whole) hold variable importance to different stakeholders and different regions is clearly an important driver of the vague use (at a global scale) of “destructive fishing”. Third, and related to the second point, we also see broad global differences in the total distribution of records (Figure 4). If a clear global use for the term “destructive fishing” is to be found, we need to ensure that the evidence base and stakeholders consulted are also global: our review suggests there is still work to be done in this area. It is particularly important to not mistake a perceived absence in one area for a complete lack of consideration; for example, Central America and West/North Africa have little discussion of destructive fishing in the media articles or academic literature analysed in this study, yet it is clearly of interest to policy-makers in these areas (Figure 4).

### Consequences of an undefined term

There remains a divide over whether to be “destructive” is to be defined by the inherent properties of a fishing gear, the case-specific nature of instances in which those gears are used, or an even wider range of parameters. For example, our study shows that the negative impacts associated with the term may include social phenomena (Fig. 5), a parameter not even considered in (FAO/UNEP, 2009).

The current debate around the role of bottom trawling in the future of wild capture fisheries exemplifies this schism and our results can partly help to explain why terminological unification could contribute to better informing this debate. In our study, bottom trawling was more associated with the term “destructive fishing” in media articles than in academic or policy documents. This is in contrast to the broader trend of media articles being less specific about gears/practices, suggesting that popular discourse drives this association more than scientific research. The only gears/practices more frequently associated with the term were “blast fishing” and “poison fishing”, which are both already politically well-established as “destructive” and, in most jurisdictions, illegal. Given we found different specificity on the use of “trawling” versus “bottom trawling” in media and policy from different continents, our results also highlight that the nuances in different terms may be understood differently in different parts of the world, and that this needs to be constructively and openly addressed.

This comparison between an already marginalized, relatively regionally-specific set of practices (“blast fishing” and “poison fishing”) and bottom trawling, a globally distributed commercial practice, exemplifies the tension between the level of evidence needed to define something as “destructive” and the politics and values associated with such a process. The question of whether bottom trawling (which is generally legal) was in the same category of “inherently destructive” as blast and poison fishing (which is generally illegal) or was “case-specifically destructive” seems to have been a major contributor to the difficulties of the previous definition-setting process (J. C. Rice, 2011). Furthermore, a 2009 review of the foundational “destructive fishing” multi-lateral framework (Figure 2) - the Code of Conduct for Responsible Fisheries (in referring to global progress on article 8.4.2 “Prohibiting destructive fishing methods and practices”) - referred to bottom trawls as “implicitly covered by the measure” but noted that very few countries have interpreted it this way and implemented full prohibitions (FAO, 2009).

The debate over the evidence and political priorities around bottom trawling remains highly polarised; several expert review studies consistently rank its environmental impact as highest amongst fishing gears (Chuenpagdee et al., 2003; Clark et al., 2017; Fuller et al., 2008). In contrast, other studies emphasise the high degree of context-specificity in ascribing “destructive” ecological effects to this practice (Hiddink et al., 2017) and the link between the severity of its impact and the level of in-situ fisheries management (Pitcher et al., 2022). While much of this debate is complex and nuanced, enduring central questions remain around whether bottom trawling is destructive in all contexts or only in specific conditions, what it means for a fishing practice to be destructive and whether there are objective parameters to identify this status. Similar problems regarding the differential interpretation and implementation of a marine policy measure have been seen in the context Marine Protected Areas and the resulting inconsistency in the protection they provide (Grorud-Colvert et al., 2021).

While a unified definition of “destructive fishing” would not resolve the intersectoral, political and value-oriented tension around the relative impacts of different fishing practices, the authors believe it would contribute strongly to better informing this debate. This in turn could foster more meaningful, consistent, and even urgent, management of cases of “destructive fishing”, in line with the requirements already established by multiple global ambitions (Figure 2).

### Recommendations for progress towards a unified definition

Our study has shown that inherent vagueness, regional variation, and deeply political schisms of interpretation may explain why global political ambitions that seek to end, prohibit or reduce “destructive fishing” have struggled to succeed. Any future process to progress towards a unified definition of “destructive fishing” and to resolve these tensions should consider the following:

1. Addressing context specificity and measurement around what is “destructive”: The tension over whether a practice is destructive at a fundamental or contextual level is the central driver of the vagueness of this term. This context includes both geographic context, and the forums (academic, media or policy) in which the term is discussed.
2. Developing a regionalised and evidence-based approach to the causal “destructive” linkages between specific fishing gears/practices and specific impacts: better capturing the interaction between gears/practices and the impacts they are associated with (across different regions) would contribute to reducing this vagueness.
3. Separating “destructive fishing” from other better-defined, fishery-associated terminology: Shared understanding is undermined where the term is elided or synonymised with other terms, for example, “overfishing” or “IUU fishing”. Separating what is “destructive” from what is merely “unsustainable” is particularly critical.
4. Recognizing (and mitigating) persistent schisms between different stakeholder groups around specific fishing practices and whether they should be considered “destructive”: The vagueness of the term also reflects long-standing and unresolved intersectoral tensions around certain practices – particularly bottom trawling and nets. Any future definition-setting process should be cognizant of these tensions and seek meaningful progress in resolving them.

The term “destructive fishing”, despite appearing in multi-lateral agreements and increasing in use over time, is used variably and vaguely across academic literature, media articles, and policy documents, as well as across geographical regions. Variation in how different stakeholder groups understand the term has no doubt contributed to tensions between cross-sectoral groups and hindered the use of “destructive fishing” in a constructive manner. Our study provides a basis of shared understanding for how the term is used in English-language documents that we hope will provide a foundation for future, constructive efforts to define “destructive fishing”.

## Acknowledgements

We are most grateful for the input from Serge Garcia, Jake Rice, Sebastian Mathew and the International Collective in Support of Fishworkers on a previous draft of the manuscript, in addition to constructive feedback from David Aldridge and three anonymous reviewers.

This project was funded by a grant from the Cambridge Conservation Initiative Collaborative Fund. D.F.W. was funded by the Department of Zoology, University of Cambridge and a Henslow Fellowship at Murray Edwards College. J.I.B. was supported by a Woolf Fisher Scholarship. D.S., J.W. and S.B. were funded by Arcadia - a charitable fund of Lisbet Rausing and Peter Baldwin.

The authors declare no conflict of interest.

## Data Availability Statement

All data is available in the manuscript or supplementary materials.

